# Muon Reduces the Training Cost of Regulatory DNA Transformers

**DOI:** 10.64898/2026.07.17.739267

**Authors:** Viraj Doshi, Malhar Bhide, Akshat Singh, Yash Rathod, Aditya Sivakumar

**Affiliations:** Origin Bio; Berkeley

## Abstract

Gene-therapy design depends on identifying regulatory sequences that drive the right level, timing, and cell-type specificity of expression. Regulatory DNA models offer a way to prioritize such sequences computationally before committing candidates to biological testing. Biological validation involves DNA synthesis, cloning, cell culture, sequencing, and functional screening, so training compute is part of the same constrained discovery pipeline rather than an isolated modeling expense. Reducing the compute required to reach a target pretraining quality could shift time and budget toward larger candidate screens, additional assays, more cell contexts, and broader follow-up validation. Given that Adam-style optimizers are widely used for training genomic sequence models, we study whether Muon can provide a more compute-efficient alternative for regulatory DNA pretraining. We provide an in-depth analysis by training Transformer models (26M–420M parameters) on ENCODE cis-regulatory sequences with Adam and Muon while holding architecture, data, and non-optimizer hyperparameters fixed and varying optimizer family, norm-control scheme, learning rate, and model width. In the largest-scale matched-target comparison, Muon reaches Adam-matched perplexity targets with a median FLOP reduction of 35.4% and a median wall-clock time reduction of 38.5%. The analysis further shows that optimizer rankings depend on norm control: independent weight decay pairs more favorably with Muon than Hyperball in this setting. These findings indicate that optimizer update structure and norm-control choices are practical levers for reducing the training resources required to reach matched perplexity targets in regulatory DNA pretraining.

## 1 Introduction

Regulatory sequence choice is a central design problem in gene therapy: the same therapeutic payload can behave very differently depending on the regulatory elements that control its expression. Models trained on regulatory DNA can help narrow this design space before biological testing, and systems such as Al-phaGenome [1], Enformer [2], and Borzoi [3] illustrate the growing role of learned sequence models in this prioritization step. Because experimental validation is costly, training compute is tied to the same constrained discovery pipeline rather than being an isolated modeling expense.

This makes optimizer choice more than an implementation detail. If one optimizer reaches the same held-out quality target with fewer FLOPs and less wall-clock time, the saved resources can support larger candidate screens, additional assays, more cell contexts, and broader follow-up validation. This study compares Adam [4] and Muon [5] for regulatory DNA Transformers under matched test perplexity targets, evaluating the optimizer in-domain rather than inferring gains from NLP. Transformer models are trained on ENCODE candidate cis-regulatory sequences while holding architecture, data, and non-optimizer hyperparameters fixed, varying optimizer family, norm-control scheme, learning rate, and model width. Adam and Muon are paired with either independent weight decay or Hyperball: **AdamW** is Adam with independent weight decay, **AdamH** is Adam with Hyperball, **MuonW** is Muon with independent weight decay, and **MuonH** is Muon with Hyperball. At the largest scale, Muon reaches Adam target quality with a median FLOP improvement of 35.4% and a median wall-clock time improvement of 38.5%. The comparison also shows that norm-control changes the optimizer ranking: independent weight decay pairs more favorably with Muon than Hyperball in this setting. These results motivate optimizer and norm-control choice as practical levers for reducing the training cost of regulatory DNA Transformers.

## 2 Related Work

Recent biological foundation model papers have started to include Muon, but mostly as a small optimizer ablation: replace Adam(W) and report the final metric. That design is useful for checking whether Muon improves a particular endpoint, but it leaves most of the optimizer behavior unmeasured. It does not show how quickly each optimizer improves during training, whether advantages appear only at some quality thresholds, whether FLOP gains agree with wall-clock gains, or whether the conclusion depends on norm control.

### Pretraining genomic language models with variants [6]

Liu *et al*. include Muon in a UKBioFormer hyperparameter ablation for gene expression prediction. Their default setting uses Adam, and replacing the optimizer with AdamW or Muon does not significantly improve performance. The comparison is useful as an endpoint ablation, but it is not designed to measure optimizer dynamics, FLOPs-to-target, wall-clock time-to-target, or interactions between optimizer family and norm control.

### Foundation models for perturbation response [7]

Cole *et al*. also report no significant difference between Adam and Muon. Their comparison tunes the learning rate, but still treats the optimizer result as an endpoint comparison. It does not factor norm control separately from optimizer family, and it does not analyze the spectral or relative step size behavior that could explain when Muon helps or fails.

### Our Contribution

We treat Muon not as a single ablation but as the subject of a detailed efficiency and dynamics study. We compare Adam and Muon across norm-control schemes, learning rates, and model widths; evaluate progression across training rather than only final loss; measure FLOPs-to-target and wall-clock time-to-target over matched quality levels; and analyze spectral entropy, participation ratio, and relative step size to understand how Muon with independent weight decay behaves differently from Muon with Hyperball. The goal is not only to ask whether Muon wins in one configuration, but to characterize when, how, and under which training dynamics it changes the cost of regulatory DNA learning.

## 3 Distribution Differences

Wilson et al. [8] showed that optimizer rankings change across domains: adaptive methods underperform SGD on some vision tasks despite converging faster, yet dominate SGD in language modeling. This is consistent with the No Free Lunch view [9] that optimizer performance depends on problem structure.

Regulatory DNA and natural language present very different sequence-learning problems. In language, much of the modeling burden comes from representing and composing meaning across a large semantic vocabulary. Regulatory DNA instead uses a compact four-letter alphabet whose predictive structure comes from biochemical sequence patterns. Regulatory signal is carried by motif identity, strength, spacing, orientation, local composition, and combinations of nearby motifs, rather than by standalone token identity. This makes regulatory DNA a distinct distributional setting for optimization: an optimizer that performs well on natural-language corpora is not guaranteed to behave the same way on regulatory sequence grammar. This study isolates that optimization question by using nucleotide cross-entropy to measure how efficiently an optimizer learns the statistical structure of regulatory DNA under matched architecture, data, and non-optimizer hyperparameters. We do not treat cross-entropy as a substitute for downstream biological evaluation on expression prediction, chromatin accessibility, or variant-effect prediction. Rather, cross-entropy probes the in-domain sequence-distribution component that such models must learn before task-specific functional heads or supervised biological readouts are added. Prior genomic foundation model results, including Evo 2 [10], suggest that lower validation loss can be associated with improved zero-shot downstream performance, but our focus here is narrower: training dynamics and compute efficiency for matched regulatory-DNA perplexity targets. Whether the same optimizer and norm-control choices improve downstream biological prediction remains an important direction for future work.

One concrete example of this distributional difference is the marginal token distribution. In language, token frequencies follow Zipf’s law [11]: a few tokens appear extremely often, while many are rare (Figure 12). This creates highly uneven token-associated gradient statistics, so Adam’s second moments can vary substantially across directions. Regulatory DNA is much less skewed. It has only four nucleotide tokens, and in our data their aggregate marginal frequencies across cell lines are nearly uniform (Figure 13), giving the corpus-level token distribution close to maximum entropy. This does not imply that individual sequences are compositionally uniform; rather, the overall token counts are far less imbalanced than in language. This potentially removes one major source of anisotropy present in language modeling, and suggests that Muon may not see the same convergence gains on regulatory DNA as it does on natural-language corpora (Appendix E).

## 4 Weight Regularization

### 4.1 Weight regularization controls optimizer step size

If the goal is to compare optimizers, the effect of weight regularization must also be controlled. Without norm control, weight norms grow during training [12], changing parameter scale relative to update size and confounding the comparison [13]. A sweep would then mix optimizer effects with uncontrolled norm drift. We therefore first use independent weight decay, which drives weight norms toward an equilibrium while separating update direction from overall parameter scale [14, 15, 12]. We choose independent rather than learning-rate-coupled decay because prior work shows stronger hyperparameter transfer for this variant [16], an important property when tuning must be done quickly and economically.

Independent weight decay is not the only plausible norm-control strategy. Although it stabilizes norms around an equilibrium, the optimizer still searches over the shell created by those fluctuations; in high dimensions, even a thin shell near the boundary can contain most of the volume [17]. For RMSNorm-sandwiched layers, scalar rescaling of a weight matrix is also partly washed out by normalization (Appendix D), making direction more important than scale. Together, these observations motivate a sharper alternative: fixing the norm and optimizing primarily over direction.

### 4.2 Hyperball as a norm control method

Hyperball [12] fixes the Frobenius norm of each weight matrix *W* to the initial radius *R* := ∥*W*_0_∥ _*F*_ and sets the norm of the update *U* to *ηR*. This confines the weight matrix to a hypersphere of radius *R*. Suppose Normalize(*A*) := *A/*∥*A*∥_*F*_ . Given an optimizer’s proposed update step *ũ*_*t*_, let the actual update be:

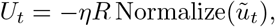

then let

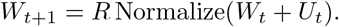

This construction can be wrapped around any optimizer update, so it works with Adam and Muon alike. That enables testing whether an optimizer advantage survives a different norm-control rule.

*Supplementary material accompanies this paper*.

The useful takeaway is that Hyperball makes the learning rate an approximate relative step size dial. This is important to control, because if the relative step size is too large, then the updates are unstable, and if it is too small, then the updates are inconsequential. Hyperball fixes ∥*W*_*t*_ ∥ = *R*, sets ∥*U*_*t*_ ∥ = *ηR*, and then retracts *W*_*t*_ + *U*_*t*_ back to the radius-*R* sphere. If *γ* is the cosine similarity between *W*_*t*_ and *U*_*t*_, in the small-learning-rate regime, this simplifies to 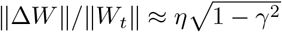 , which becomes ≈*η* when the update is nearly orthogonal to the current weights (Appendix B); a corresponding bound is given in Appendix C.

#### Relative step size of weight decay

Under weight decay, by contrast, the relative step size depends on the learning rate, decay coefficient, optimizer state, momentum coefficients, and training time [14, 12, 18]; Hyperball removes most of that coupling by fixing ∥*W*_*t*_∥ = *R* and ∥*U*_*t*_∥ = *ηR*. In our runs, MuonW’s relative step size decreases as ∥*W*_*t*_∥ grows, while MuonH remains flatter at the level set by *η* (Figures 10 and 15).

## 5 Experimental Setup

To isolate optimizer effects, we hold the architecture family, dataset, processed-token budget, input pipeline, output heads, and all non-optimizer hyperparameters fixed. We vary only optimizer family, norm-control scheme, learning rate, and model width. All runs use weight decay 10^*−*5^ and train for 20 epochs on ENCODE v4 [19] candidate cis-regulatory elements specific to one of eight cell lines (Appendix A). The held-out set consists of distinct regulatory sequences not used for training. The eight-cell-line subset is small enough that training necessarily revisits examples across epochs, but large enough that each epoch contains many distinct regulatory sequences. This keeps the comparison compute-bounded rather than data-bounded: optimizer differences are measured by how efficiently each method uses a fixed data distribution over repeated passes. We test three Transformer widths: 512, 1024, and 2048, corresponding to approximately 26M, 105M, and 420M parameters. Runs use 8-GPU NVIDIA H100 80GB nodes. Inputs are tokenized at single-nucleotide resolution and trained with a masked language modeling objective. The nucleotide head provides the held-out cross-entropy and perplexity metrics, while the modality head provides an auxiliary DNase-seq prediction loss using a factorized Poisson and multinomial objective. Figure 1 summarizes the model and training objectives.

**Figure 1:**
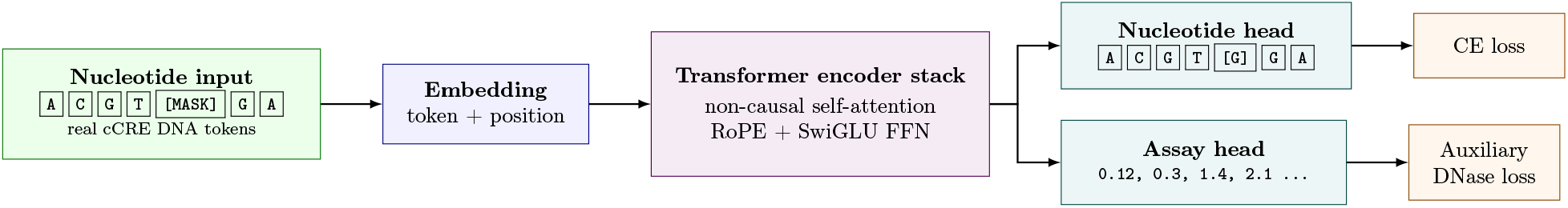
Model Architecture.

## 6 Results

### 6.1 Muon reaches Adam perplexity targets with median 35.4% fewer FLOPs and 38.5% less wall-clock time

Lower perplexity indicates better probability mass placement on the target DNA sequence. We convert checkpoints to FLOPs using a base forward/backward term proportional to 6*PBL* per step plus a per-optimizer update term; in the largest setting, Muon’s Newton–Schulz update adds only about 0.94% overhead relative to the base forward/backward pass (Appendix F).

We first aggregate optimizer variants by family to show the high-level efficiency trend. In Figure 2, each run is trained for 20 epochs and checkpointed once per epoch. At each checkpoint, the family curve reports the best held-out perplexity achieved by any run in that optimizer family, mapped back to its FLOP and wall-clock cost. Across the relevant compute and time range, the Muon envelope stays below the Adam envelope, showing that Muon reaches better held-out perplexity over a broad set of budgets rather than only at a single endpoint.

**Figure 2:**
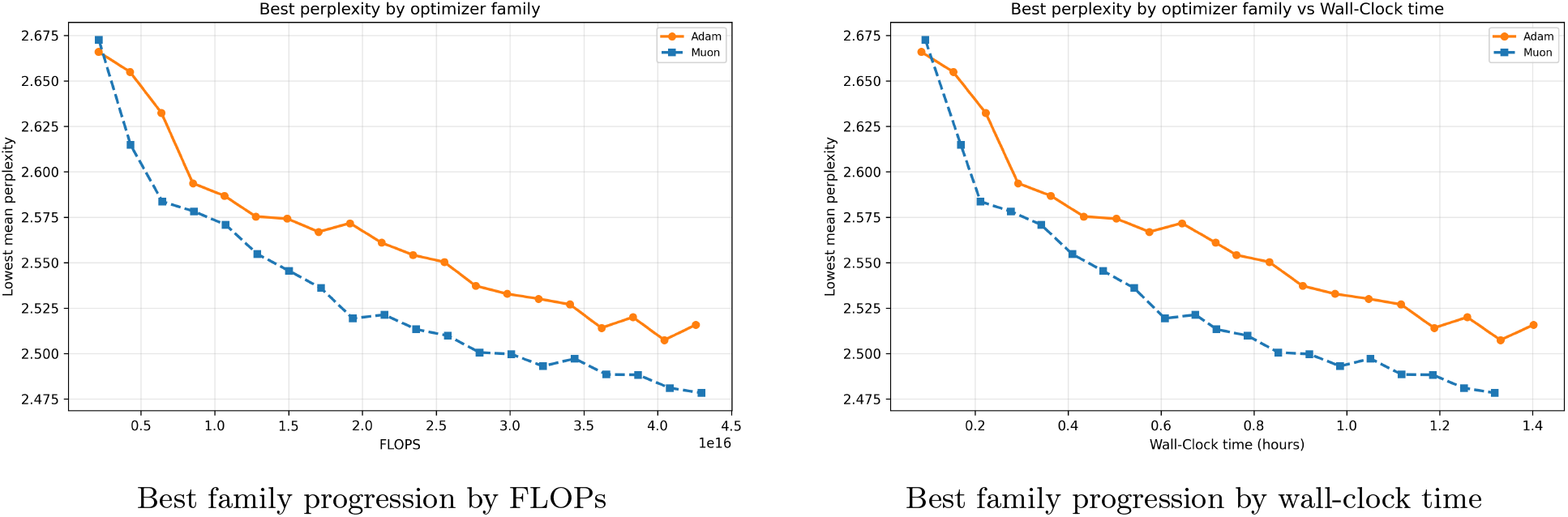
Largest-scale family-level progression efficiency.

Figure 3 turns these progression curves into a matched-quality efficiency comparison. For each Adam checkpoint, we use its held-out perplexity as the target and find the lowest-cost Muon checkpoint whose perplexity is at least as good. The plotted value is the resulting FLOP or wall-clock time reduction relative to that Adam checkpoint, with positive values indicating that Muon reaches the same target perplexity with less compute or time. After the earliest targets, the Muon advantage remains consistently positive and typically falls in the 35–45% range for both FLOPs and wall-clock time.

**Figure 3:**
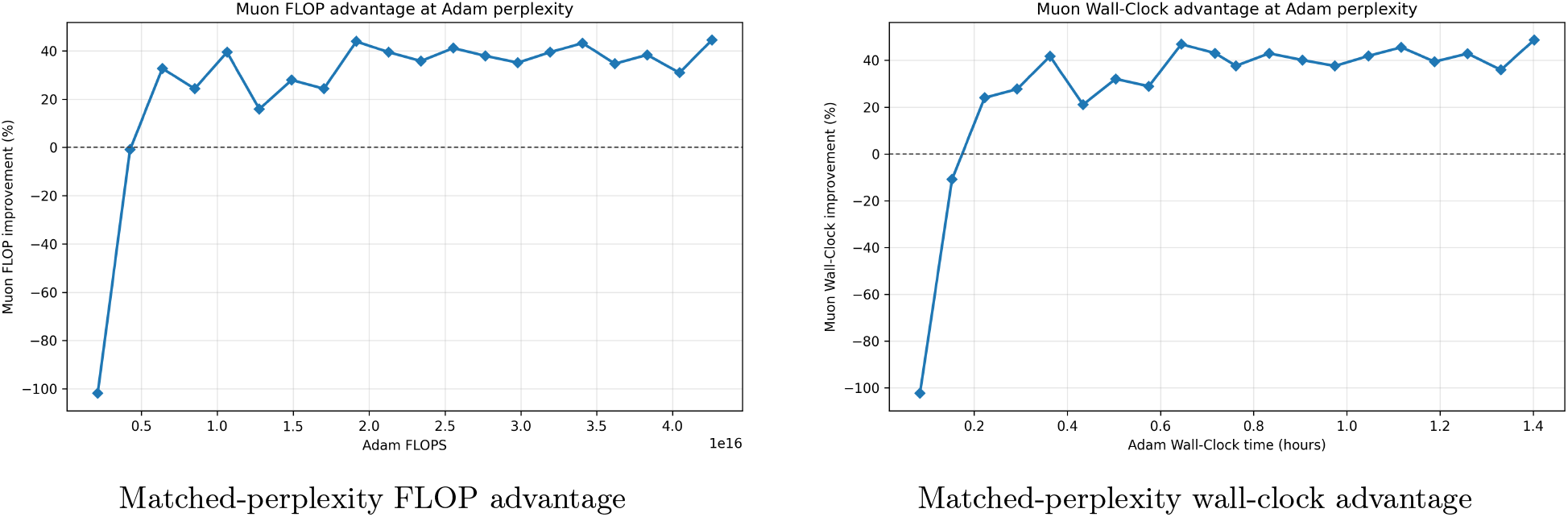
Muon advantage at matched Adam perplexity.

Figure 4 summarizes the matched-target comparison across tested model scales. For each scale, we report the median Muon reduction across Adam perplexity targets. The median advantage is positive across the tested scales: 18.3–39.7% fewer FLOPs and 24.6–42.0% less wall-clock time, suggesting that Muon’s efficiency gain is not confined to one model width.

**Figure 4:**
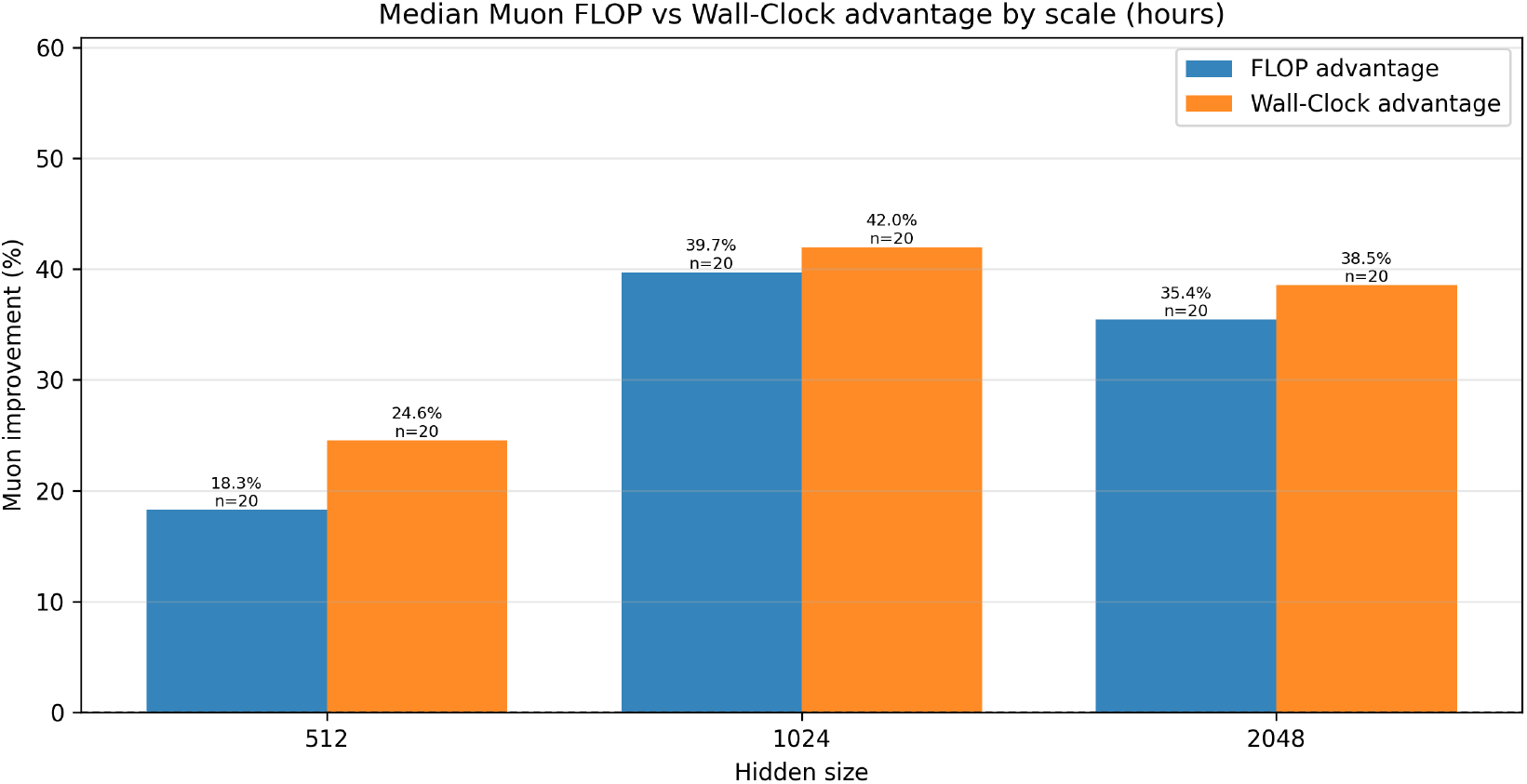
Median Muon FLOP and wall-clock advantage across matched Adam perplexity targets by scale.

### 6.2 Test perplexity across sweep grid

Figure 5 gives an optimizer-level view at the largest scale, using the run with the lowest final EMA perplexity for each optimizer variant; the EMA values used for this selection are tabulated in Appendix G. This complements the matched-target wall-clock analysis above by showing mean test perplexity as a function of training FLOPs. The curves separate clearly: SGD remains well above the Adam and Muon variants, Adam improves steadily, and the best Muon configuration reaches its lowest perplexity on a smaller compute and wall-clock budget. The selected Muon run uses learning rate (LR) 2.27 *×* 10^*−*2^, almost an order of magnitude larger than the LR 5.69 *×* 10^*−*3^ for the strongest Adam run, suggesting that Muon’s update structure is associated with a larger stable effective step size in this sweep, consistent with faster progress toward the same held-out perplexity target.

**Figure 5:**
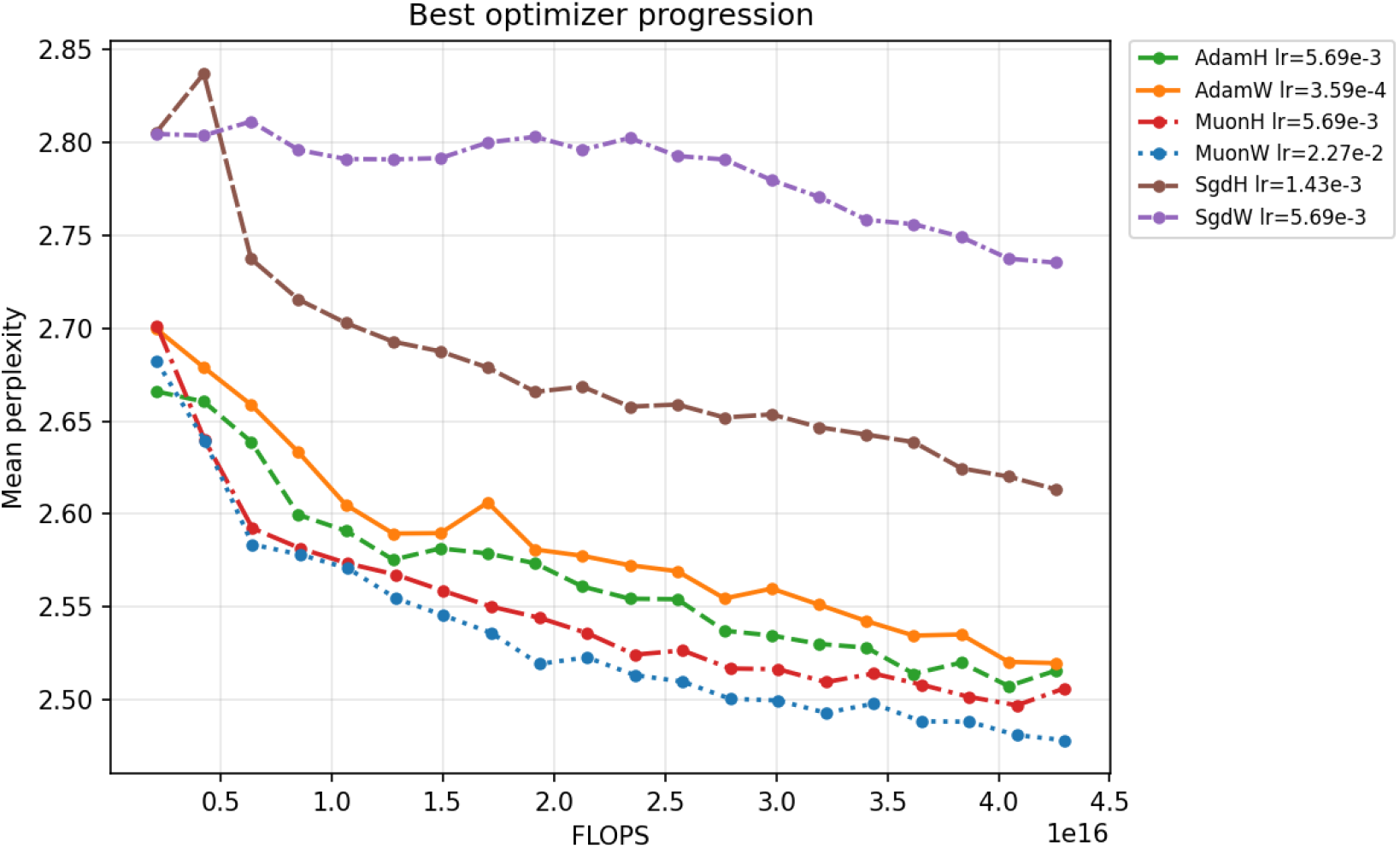
Largest-scale (2048) optimizer progression.

### 6.3 Independent weight decay is more favorable for Muon than Hyperball

In this sweep, MuonW is the strongest configuration, showing that Muon’s advantage depends on the norm-control scheme as well as the update rule itself. This raises the question of whether MuonW and MuonH differ only in their instantaneous updates, or also in how those updates accumulate into the weights. Section 8 tests this emperically and suggests that independent weight decay better preserves Muon’s effective rank through training, while Hyperball yields a more concentrated learned weight spectrum.

## 7 Training Dynamics

The matched-target results show that Muon reaches lower held-out perplexity levels with less compute and time, but they do not explain the source of this advantage. To probe the mechanism associated with this advantage, we analyze training loss sweeps for the within-optimizer norm-control comparisons in Figures 6 and 7, the strongest cross-optimizer comparison in Figure 8, and aggregate learning curves for the perplexity-selected runs in Figure 14.

**Figure 6:**
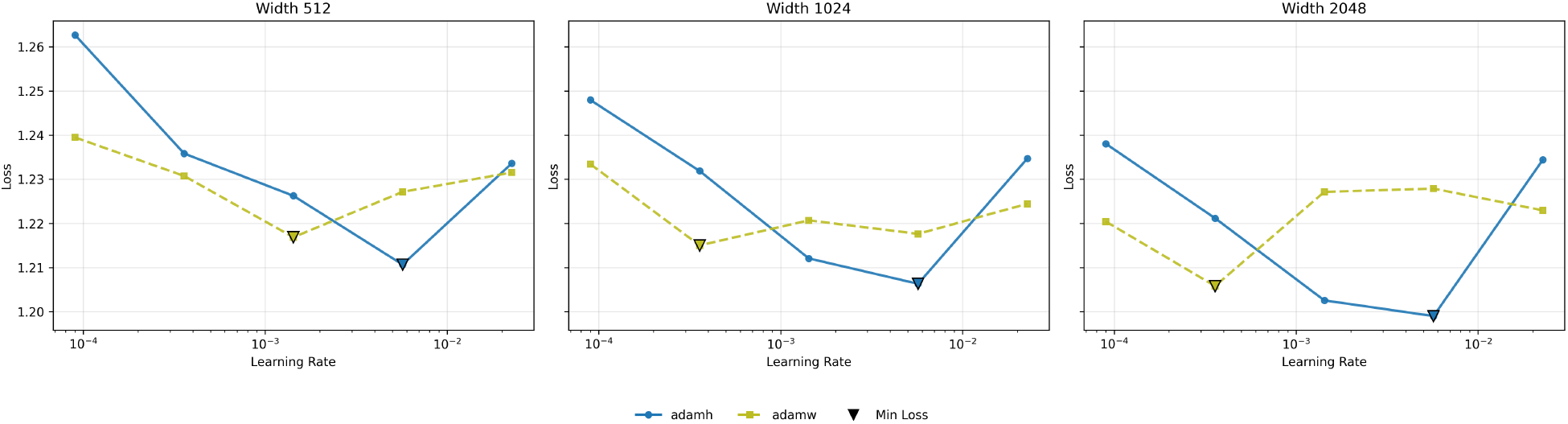
AdamW–AdamH training loss sweep.

**Figure 7:**
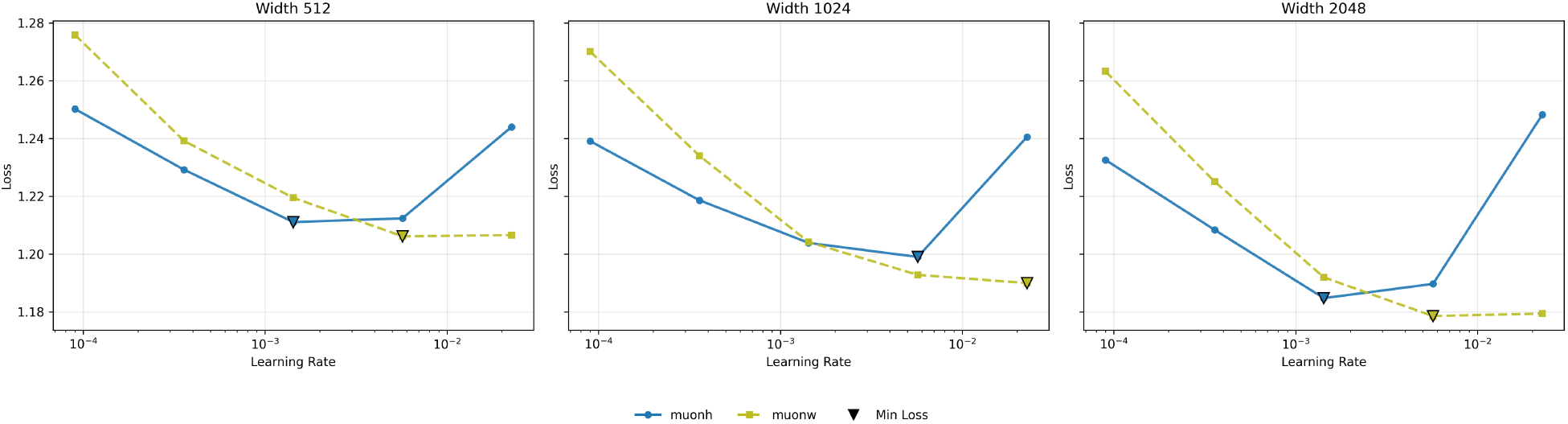
MuonW–MuonH training loss sweep.

**Figure 8:**
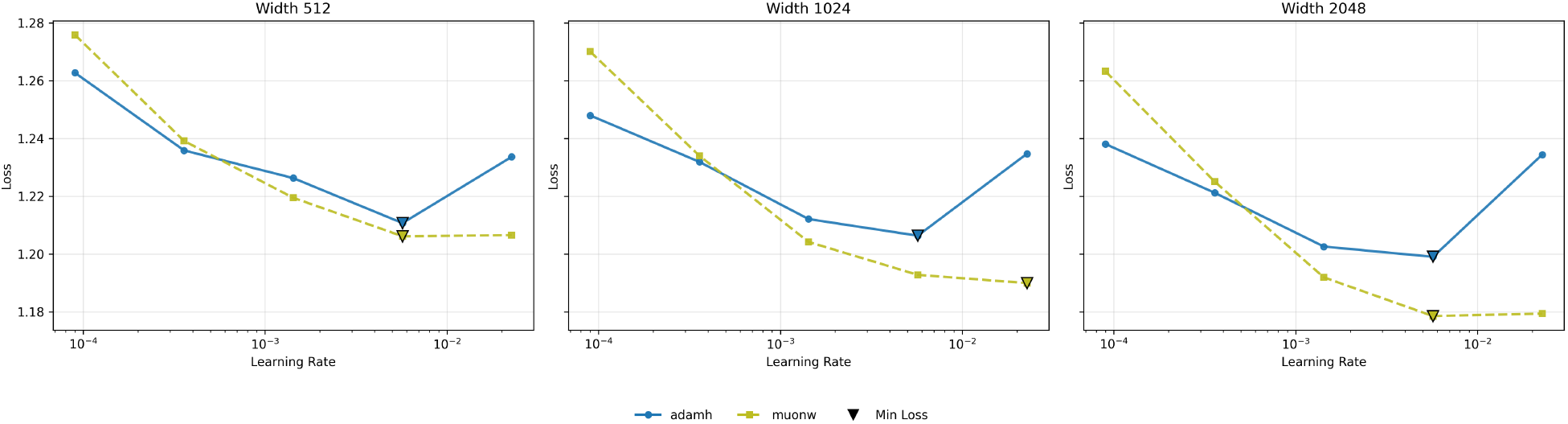
MuonW–AdamH training loss sweep.

Consistent with prior work [5, 12], Muon reaches lower training loss than Adam, and the optimizer ranking matches the main result: MuonW converges fastest, followed by MuonH, AdamH, and AdamW. The central pattern is learning-rate dependence. AdamH is most competitive in the mid-range, AdamW is more stable at the extremes, MuonH performs best at smaller learning rates, and MuonW improves most as the learning rate increases. Although the exact training-loss optimum for MuonW varies across widths, LR ∼5.69 *×* 10^*−*3^ for widths 512 and 2048, versus LR ∼2.27 *×* 10^*−*2^ for width 1024, the broader trend is that MuonW pulls away from AdamH most clearly in the high-learning-rate regime, especially at larger width. This supports the step-size hypothesis from Figure 5: MuonW’s gains arise from its ability to sustain a larger stable effective step size, making higher learning rates increasingly useful at scale.

## 8 Spectral Analysis

To characterize differences between Muon with independent weight decay (MuonW) and its constrained counterpart (MuonH) in this sweep, we examine the spectral properties of the learned representations. Following the eigenspectral framework of NerVE [20]^1^, we track two key metrics for the weight matrices and the corresponding parameter updates:

- **Spectral Entropy (SE):** A measure of how uniformly variance is distributed across singular values. Values near 1.0 indicate a broadly distributed spectrum, while lower values indicate stronger spectral concentration. This metric is scale-independent as it is computed after normalizing the spectrum.
- **Participation Ratio (PR):** The effective rank of the matrix. We normalize the PR to get the ratio of the effective rank to full rank. This metric is scale independent as it is the ratio of the square of the sum of eigenvalues to the sum of the squares of the eigenvalues of the gram matrix.

Figure 9 shows that Hyperball narrows Muon’s accumulated weight spectrum without fully removing its update diversity. Both Muon variants start with weight spectral entropy near 0.98, but by the end MuonW remains broader than MuonH: SE is approximately 0.95 versus 0.93, and normalized PR is approximately 0.57 versus 0.46. AdamW reaches similar SE to MuonW but with lower PR, approximately 0.36, while AdamH is the most concentrated configuration, with SE near 0.87 and PR near 0.14. This exposes a deceptive aspect of SE: a spectrum can look high-entropy while still placing too little mass in enough useful directions. In this figure, PR separates the successful configuration more cleanly than SE, suggesting that lower held-out perplexity aligns more closely with maintaining effective rank than with maximizing entropy alone.

**Figure 9:**
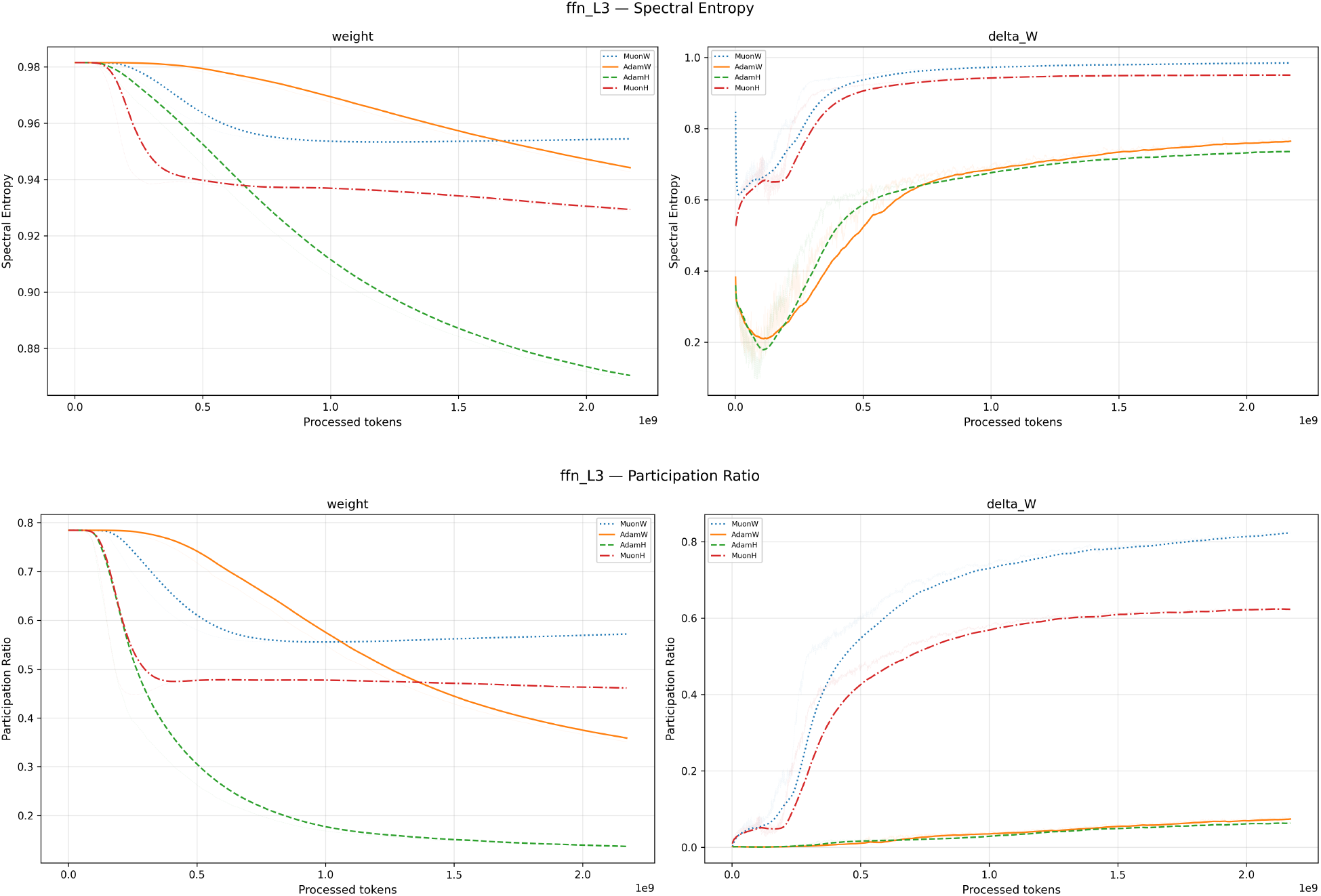
Spectral metrics for FFN Layer 3. Top: Spectral Entropy. Bottom: Participation Ratio. Each row shows the spectrum of the weights on the left and the mean update Δ*W* on the right. **Blue:** MuonW, **Red:** MuonH, **Orange:** AdamW, **Green:** AdamH.

**Figure 10:**
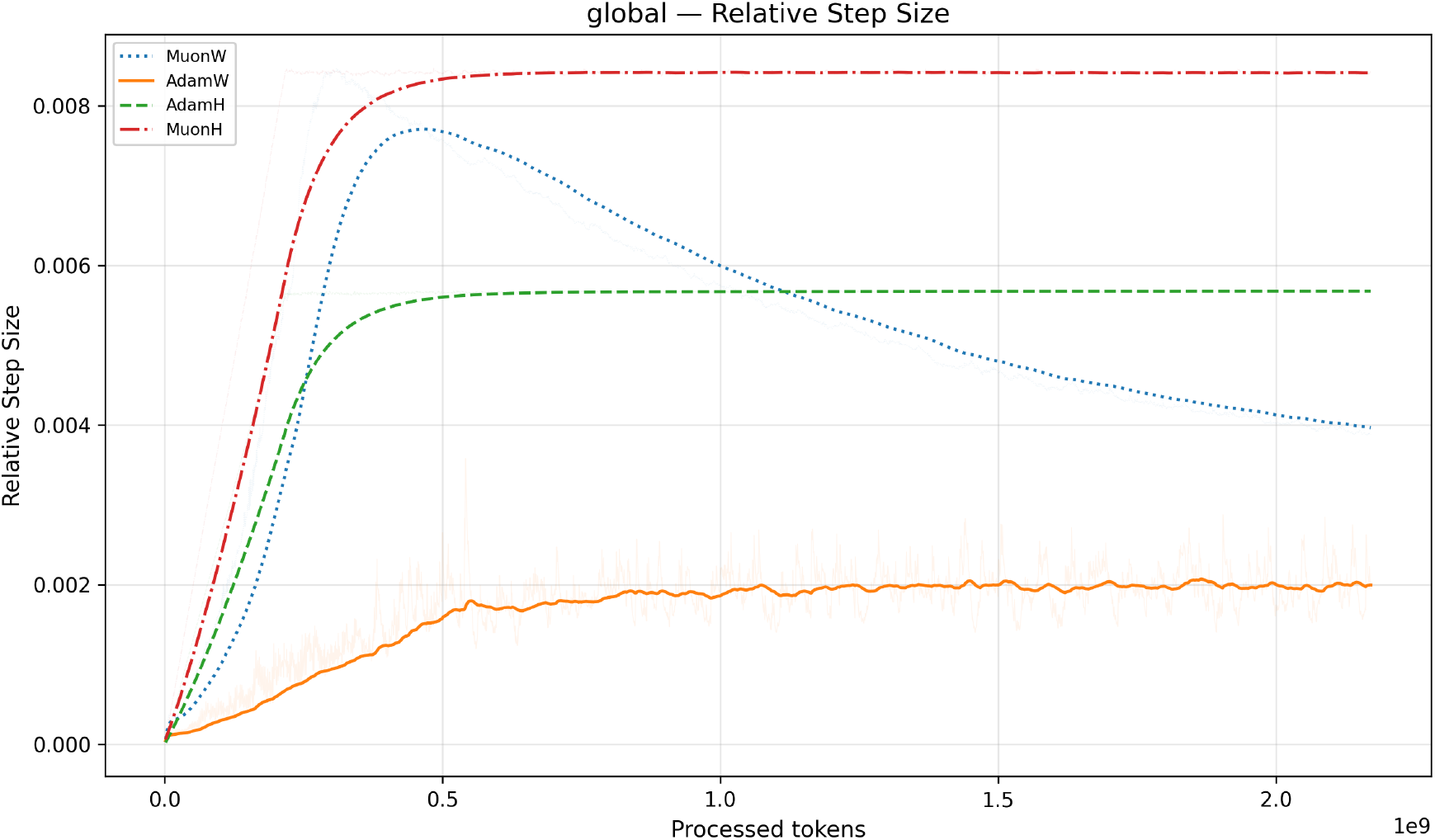
Relative step size.

**Figure 11:**
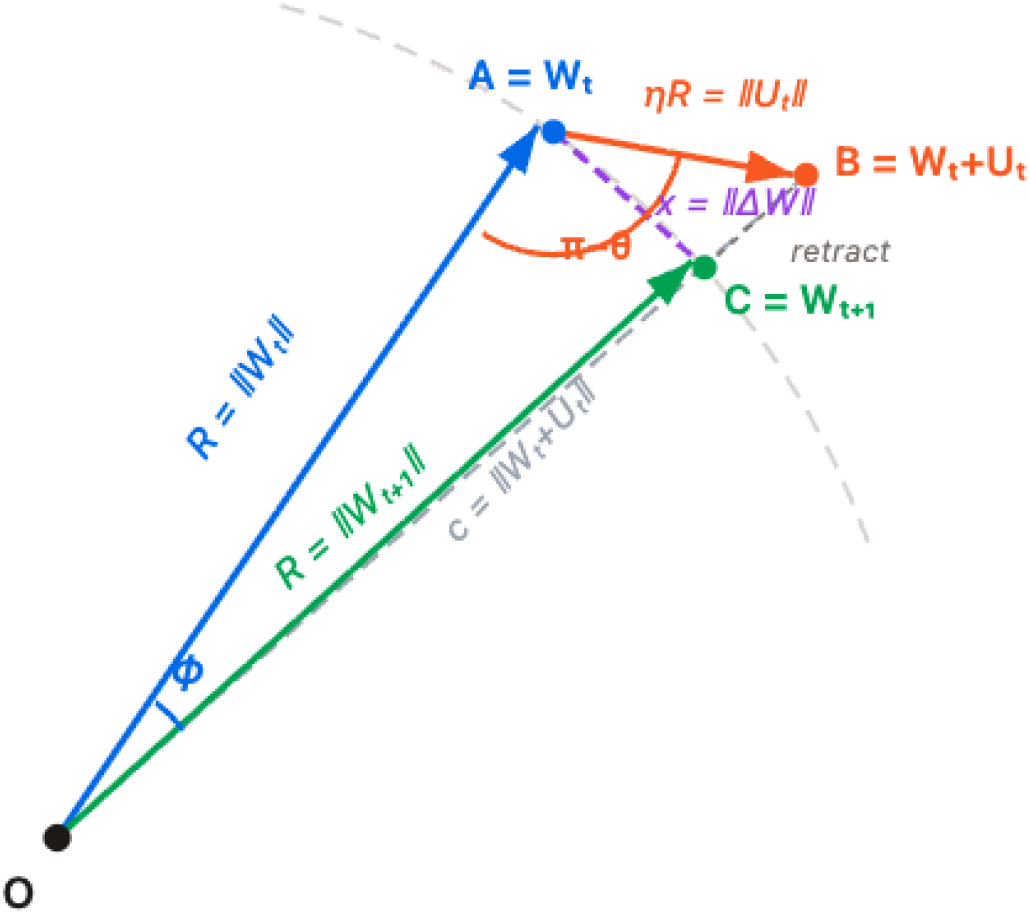
Hyperball retraction geometry.

**Figure 12:**
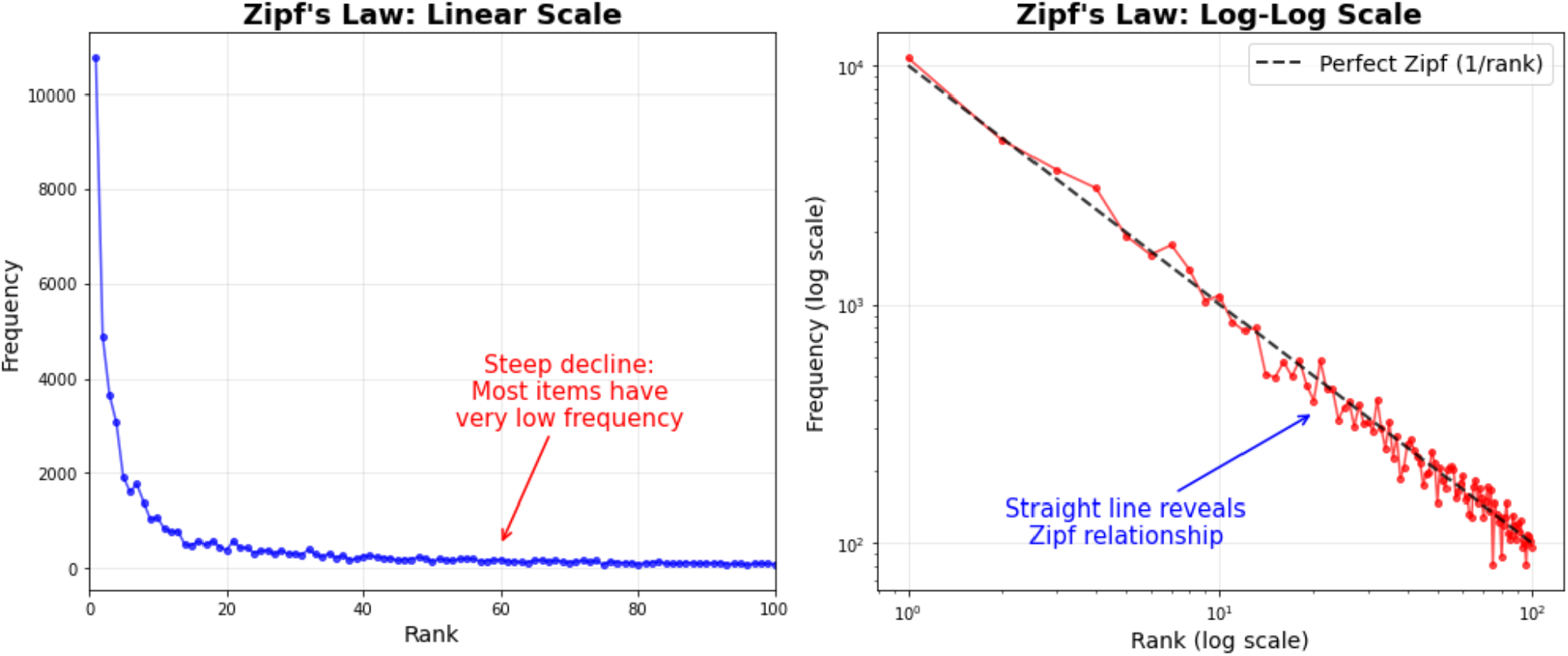
Zipf’s law. The left panel shows Zipf’s law on a linear scale, and the right panel shows it on a log-log scale. The straight line on the log-log plot is the signature of a power-law distribution [23].

**Figure 13:**
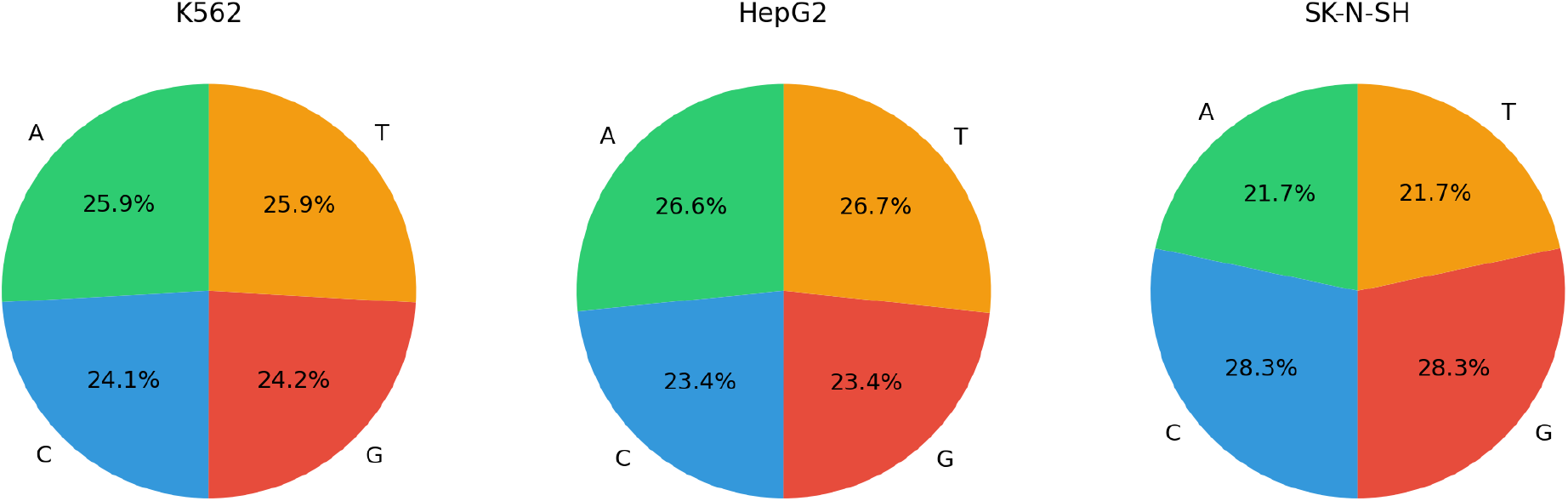
Nucleotide frequencies. The panels show nucleotide frequencies across three random cell lines in our training set.

**Figure 14:**
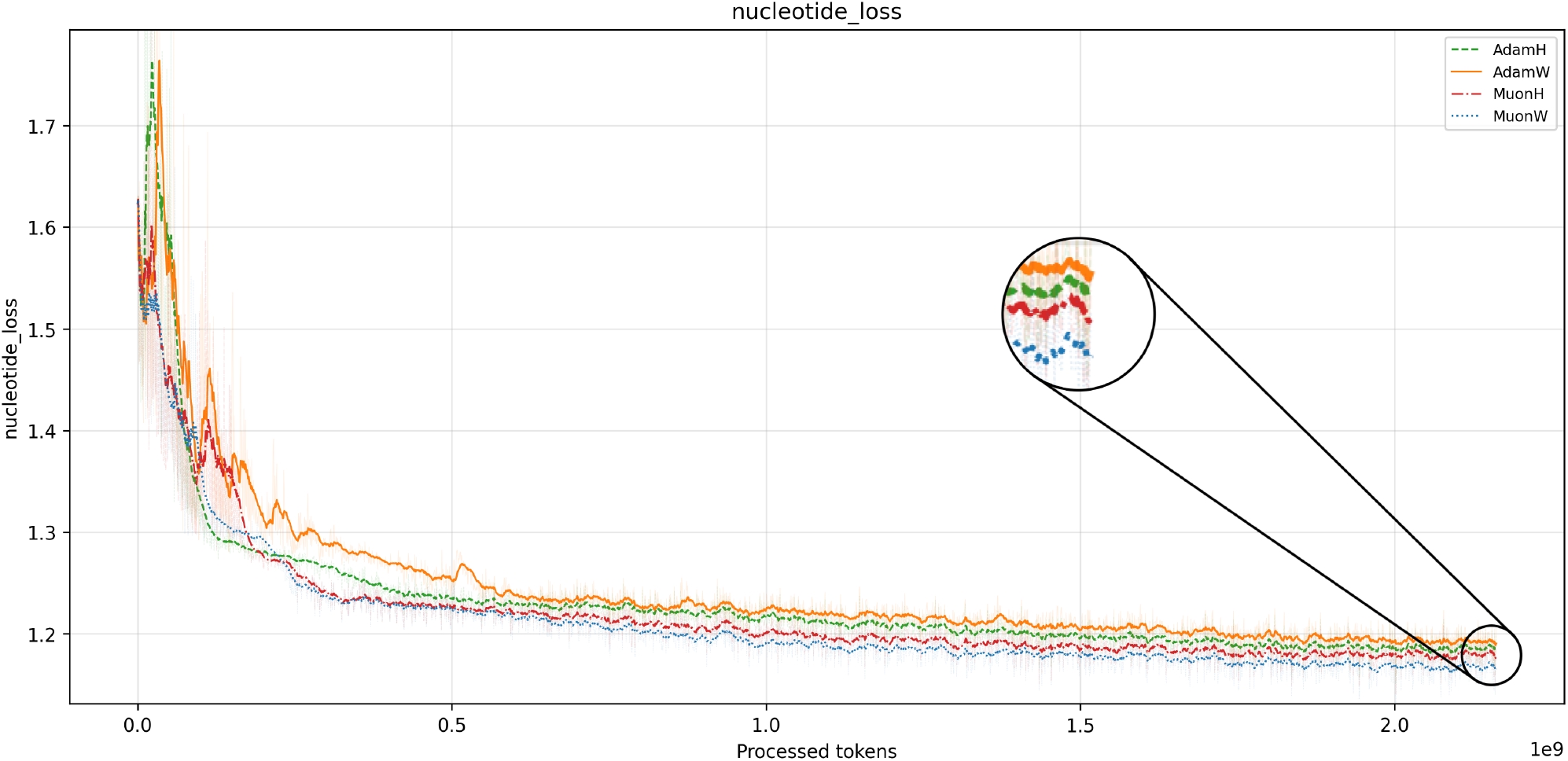
Aggregate training loss curves for the perplexity-selected runs across optimizer combinations.

**Figure 15:**
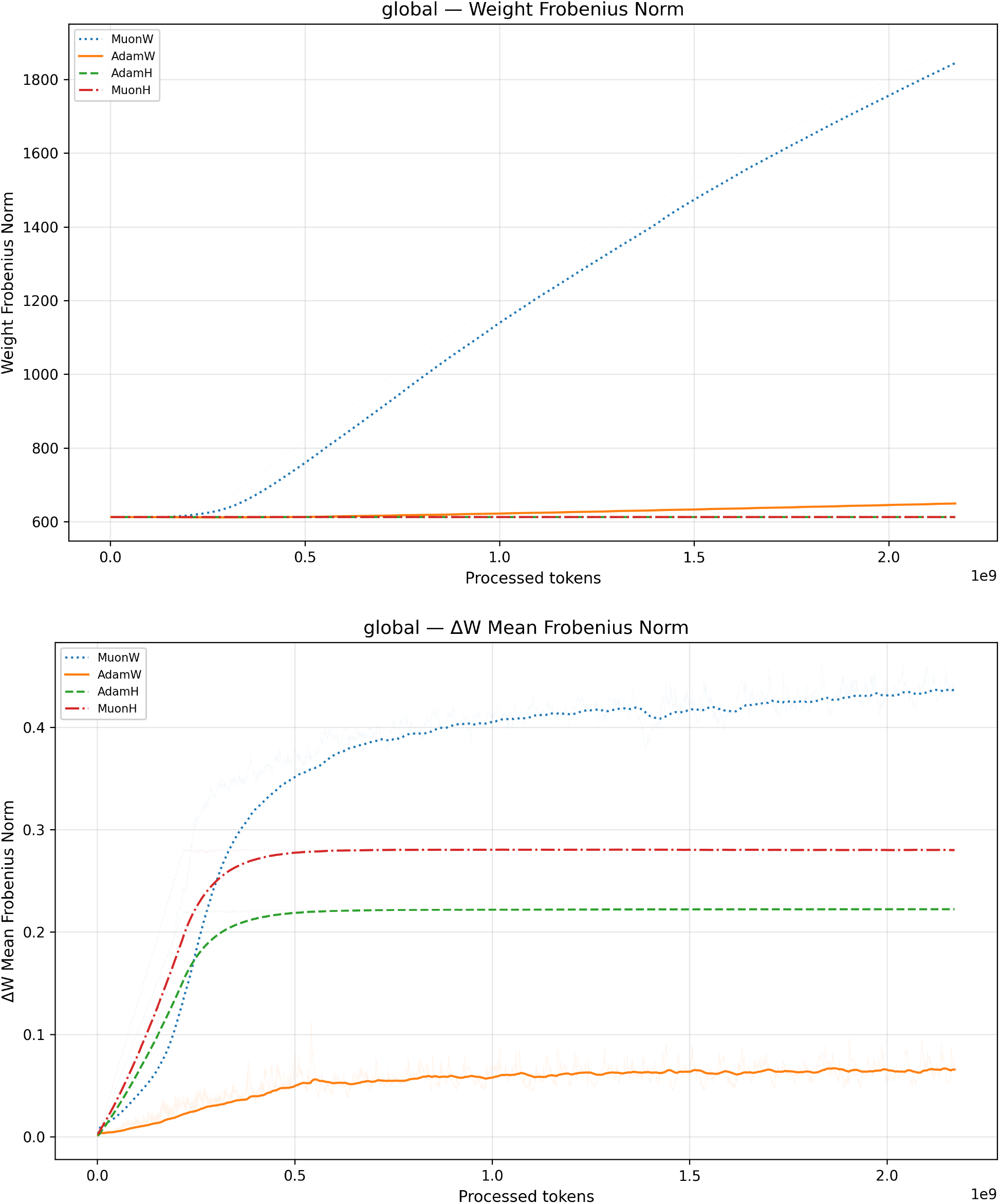
Weight and update norms. The top panel shows weight Frobenius norm, and the bottom panel shows update mean Frobenius norm. MuonW shows monotonically increasing weight norm with roughly constant update norm.

### 8.1 Uniform Rescaling and Spectral Concentration

Figure 9 separates two questions: what spectrum the optimizer produces in its instantaneous updates Δ*W* , and what spectrum ultimately accumulates in the weights *W* . Both Muon variants produce much broader update spectra than Adam: their update PR is far higher, meaning their steps use many more independent matrix directions. However, this update diversity is not transferred equally into the learned weights. MuonW maintains the broadest weight spectrum, while MuonH ends with lower weight SE and PR. This suggests that Hyperball does not remove Muon’s high-rank update structure at each step; instead, its repeated Frobenius retractions appear to alter the training trajectory and limit how much the weaker singular directions can accumulate in *W* . Because SE and PR are scale-independent, this difference is not a trivial consequence of rescaling the matrix norm; it reflects a different optimization trajectory under Hyperball. AdamW shows why PR is important: it keeps relatively high weight SE, but its lower PR indicates that the usable effective rank remains smaller than MuonW’s. Overall, the configuration with the best held-out perplexity, MuonW, is also the one that best preserves effective rank from updates into weights.

### 8.2 Implicit Annealing and Relative Step Size

Figure 10 suggests a second reason MuonW outperforms MuonH: its relative step size, ∥Δ*W*∥ */* ∥*W*∥ , naturally decays over training. MuonW’s weight norm grows while its update norm stays roughly stable (Figure 15), so each update becomes smaller relative to the current weights. This resembles an implicit annealing effect: training begins with larger relative moves and later shifts toward finer adjustments. MuonH lacks this pattern because Hyperball fixes the weight norm, causing its relative step size to plateau. AdamW shows weaker norm-growth annealing because its weight norm grows much more slowly, and its coordinate-wise updates lack Muon’s matrix-whitened structure: Adam’s noisy per-parameter changes can cancel in the first moment *m*_*t*_ while continuing to accumulate in the second moment *v*_*t*_, shrinking 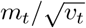 in noisier coordinates. Overall, MuonW appears to combine high-rank coherent updates with automatic late-training step-size refinement, while MuonH preserves the coherent-update side but not the same relative-step decay.

## 9 Discussion and Limitations

This work shows that optimizer and norm-control choices can materially change the training cost of regulatory DNA Transformers under a fixed architecture, dataset, and objective. MuonW reaches matched Adam perplexity targets with fewer FLOPs and less wall-clock time, and the spectral analysis provides a possible explanation: the winning configuration is also the one that best preserves effective rank through training. More broadly, these findings support treating regulatory DNA pretraining as its own optimization setting rather than assuming that optimizer behavior from language modeling transfers unchanged [8, 9]. These findings should be read with several limitations in mind:

- **Downstream biology:** Held-out perplexity is the matched target. We do not show improved expression prediction, chromatin-accessibility prediction, variant-effect prediction, or regulatory sequence design.
- **Statistical variation:** The sweep used 90 runs of 20 epochs, but did not include multiple seeds for each setting. We therefore do not report error bars or statistical significance tests.
- **Fixed training recipe:** Batch size, sequence length, masking setup, architecture, data mixture, and weight decay (10^*−*5^) were held fixed to isolate optimizer effects. The ranking may change under other recipes.
- **Scale:** The experiments cover models up to the few-hundred-million-parameter regime. Optimizer behavior may differ at larger scales.

## Appendix

### A Data

We construct the dataset from the ENCODE v4 candidate cis-regulatory element data [19]. We preprocess data exactly as described in the paper, except for dividing by the sample non-zero mean during batch creation. Starting from the ENCODE v4 cCRE set, we filter for examples whose DNase-seq activity is specific to exactly one of the eight cell lines used in our experiments (A549, HEK293T, HepG2, Jurkat Clone E6-1, K562, SK-N-SH, endothelial cell of umbilical vein, skeletal muscle myoblast). This yields regulatory sequences of length ≤ 350 with a single active DNase track among the selected cell contexts, which supports the auxiliary modality-prediction objective while keeping the training distribution focused on cell-type-specific regulatory activity.

#### Assets and licenses

We use ENCODE v4 candidate cis-regulatory element annotations and AlphaGenome preprocessing resources. ENCODE data are released for unrestricted use under the ENCODE data release policy, and the Google DeepMind AlphaGenome code resources are released under the Apache License 2.0. We cite the original sources and follow their stated terms of use. No new dataset, benchmark, or model checkpoint is released as part of this submission. The subset is available at https://kaggle.com/datasets/bee59cfcf029b7fa33440aebf35e01a8039aa27982de870e323b27afe626899e.

### B Full Derivation of the Relative Step Size

The Hyperball retraction maps *W*_*t*_ + *U*_*t*_ back onto the hypersphere of radius *R*. The geometry is captured by two triangles in the vectorized matrix space:

1. **Triangle** *O***-***A***-***B*: where *O* is the origin, *A* = *W*_*t*_ lies on the sphere, and *B* = *W*_*t*_ + *U*_*t*_ is the pre-retraction sum (generally off the sphere). Sides: ∥*OA*∥ = *R*, ∥*AB*∥ = *ηR*, ∥*OB*∥ = *c*. Interior angle at *A*: *π − θ*. angle *ϕ*.
2. **Triangle** *O***-***A***-***C*: where *C* = *W*_*t*+1_ = *R* Normalize(*W*_*t*_ + *U*_*t*_) is the retracted weight, also on the sphere. This is an isosceles triangle with ∥*OA*∥ = ∥*OC*∥ = *R*, chord ∥*AC*∥ = *x* = ∥Δ*W* ∥, and apex

Figure 11 shows the 2D cross-section used for the derivation, with the relevant lengths and angles labeled. We now derive the exact step size. Using the cosine rule on triangle *O*-*A*-*B* (where *O* is the origin, *A* = *W*_*t*_, and *B* = *W*_*t*_ + *U*_*t*_), noting that the interior angle at *A* is *π − θ*:

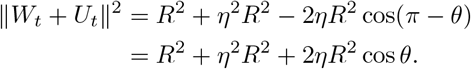

Hence the pre-retraction length is 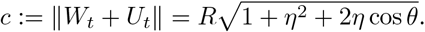 .

Since *W*_*t*+1_ lies on the ray from *O* through *B*, the angular change 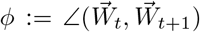 equals ∠*AOB*. Applying the sine rule to △*OAB*:

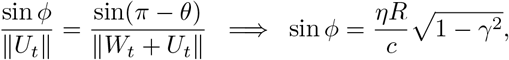

and therefore

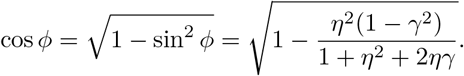

Finally, *W*_*t*_ and *W*_*t*+1_ both lie on the sphere of radius *R*, so we apply the cosine rule to the isosceles triangle △*OAC*:

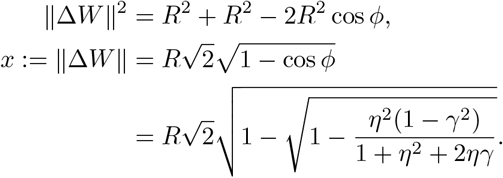

This exact expression also recovers the intuitive *O*(*η*) relative step size argument. Hyperball sets both ∥*W*_*t*_∥ = *R* and ∥*U*_*t*_∥ = *ηR*. If ∥Δ*W* ∥_*F*_ is on the same order as ∥*U*_*t*_ ∥_*F*_ , then

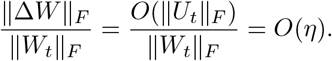

The exact formula above refines this by accounting for the alignment between *W*_*t*_ and *U*_*t*_.

#### Small-learning-rate regime

In our practical training regime, *η*^2^ ≪ 1 and *γη* ≪ 1, so 1 + *η*^2^ + 2*ηγ* ≈ 1 and

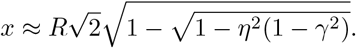

Applying 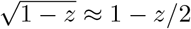 yields

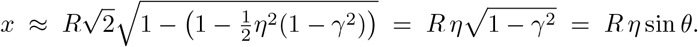

Since ∥*W*_*t*_∥ = *R*, the relative step size is

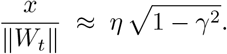

When the update is orthogonal to the current weights (*γ* = 0), this collapses to 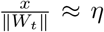 . Thus, in this regime, doubling the learning rate approximately doubles the effective movement on the sphere.

### C Bounds on the Relative Step Size

We restrict to *γ, η* ∈ (0, 1) throughout this appendix; the *γ* ∈ (*−*1, 0) case follows by replacing *γ* → |*γ*| in every step (the formula is dominated by *γ*^2^, and the linear 2*ηγ* term in the denominator only tightens the bound when *γ <* 0). Now, suppose we know that *γ, η* ∈ (0, 1). We can infer from this that:

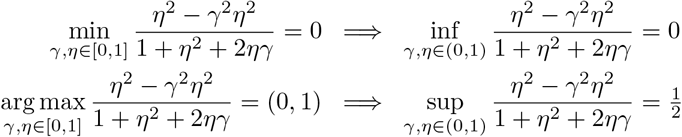

From this we obtain

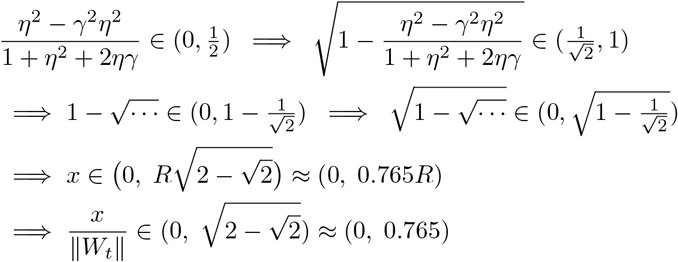

This is the tightest bound on the relative step size obtainable without knowledge of *γ*. Computing *γ* analytically remains an open question, as the repeated retractions onto the hypersphere make it difficult to translate previous arguments involving momentum and weight decay [12]. Unfortunately, this bound is not particularly informative: the lower bound is vacuous, and the upper bound is substantially looser than the ideal *O*(*η*). Now, if, for instance, we input a standard range ∼*η*∈ (10^*−*4^, 10^*−*1^) and fix *γ* = 0, we get a slightly tighter bound of (0.0001, 0.0996), where a higher learning rate corresponds to a higher step size. For a higher dot product like *γ* = 0.5, we get a bound of (0.0000866, 0.0823), which is lower.

### D Enforcing Scale Invariance

A key ingredient is RMSNorm [21], which rescales a vector by its root-mean-square (RMS) magnitude. Given an activation vector *x* ∈ ℝ^*d*^, RMSNorm computes

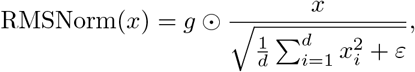

where *g* ∈ ℝ^*d*^ is a learned per-coordinate gain, *ε* > 0 is for numerical stability, and ⊙ is the Hadamard product (element-wise multiplication).

#### Rescaling property

Ignoring *ε* (or when ∥*x*∥ is not tiny), RMSNorm is *scale invariant* : for any scalar *α* > 0,

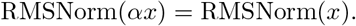

Intuitively, scaling the input by *α* scales the denominator by the same *α*, so the normalized direction stays the same.

#### Scale invariance of weight matrices

Consider a linear map *y* = *W x*. If the model applies RMSNorm before the next computation (pre-norm, post-norm, or both), then multiplying the weight matrix by a scalar often has little effect on the downstream activations:

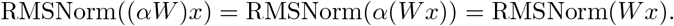

So for these “RMSNorm sandwiched” layers, scale doesn’t matter as the RMSNorm layer itself learns the scaling factor necessary for the output.

In our model architecture, we used MuP initialization and layer pre-normalization, but that was not enough to enforce scale invariance. To further enforce it, we added a layer post-norm and then QK-norm (like Gemma 3’s and 4’s architecture [22]), which appeared to be sufficient for our model (although adding only the post-norm may already have been enough; the QK-norm was added for completeness).

### E Gradient Statistics Differ Between DNA and NLP

Consider a vocabulary of *K* tokens with marginal frequencies *p*_1_ ≥ *p*_2_ ≥ *· · ·*≥*p*_*K*_. In a mini-batch of *B* tokens drawn i.i.d. from *p*, the expected count of token *k* is *n*_*k*_ = *B p*_*k*_. The gradient of the loss decomposes as

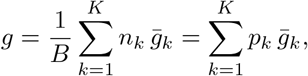

where 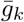 is the mean gradient contribution conditioned on token *k*. We can say that Adam’s running second moment for parameters associated with token *k* accumulates as 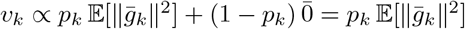 . The ratio of the largest to smallest second moment therefore scales with the frequency ratio of the most to least common token 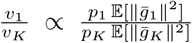 . If we assume that 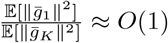 , then:

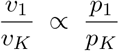

#### Zipf (NLP)

With *p*_k_ ∝ 1*/k* and *K* ≈ 100,000, the dynamic range is *p /p* ≈ *K* ≈ 10^5^. Adam’s per-coordinate 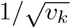 scaling spans *orders of magnitude*, providing substantial per-direction adaptation that compensates for the wildly uneven gradient contributions of frequent vs. rare tokens.

#### DNA

With *K* = 4 and *p*_*k*_ ≈ 1*/*4, the dynamic range is *p*_1_*/p*_*K*_ ≈ 1. Adam’s adaptive denominator 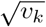 is nearly identical across all tokens, reducing the coordinate-wise scaling to an approximately uniform scalar. The per-direction adaptation that makes Adam powerful in NLP provides *negligible benefit* here, while Muon’s whitened, scale-free updates are naturally suited to this near-isotropic landscape.

### F FLOPs Accounting

Each FLOPs-axis plot uses a per-optimizer cost model that accounts for both the per-token forward/backward compute and the per-step optimizer update. Let

- *L*: sequence length (tokens per sequence),
- *B*: batch size (sequences per step),
- *N*: total sequences in the dataset,
- *P*_2D_: total 2D parameters (linear weights and embeddings); optimized by Muon/MuonH in mixed mode and dominating forward/backward matmul compute,
- *P*_non-2D_: total non-2D parameters (RMSNorm scales, any biases); under the mixed-Muon optimizers (MuonW and MuonH) these are handled by an AdamW sub-optimizer, while under plain AdamW or AdamH all parameters share the top-level optimizer,
- *d*: hidden dimension,
- *C*_opt_: per-parameter FLOP cost of a single optimizer step.

Total FLOPs for one epoch decompose into a *base compute* term (forward/backward, incurred once per token) and an *optimizer overhead* term (incurred once per batch step). We use the standard 6 *P*_2D_-FLOPs-per-token matmul proxy for the base compute, which ignores non-matmul ops (attention softmax/mask, activations, norms, embedding lookups) contributing a few percent at *L* = 352, *d* = 2048:

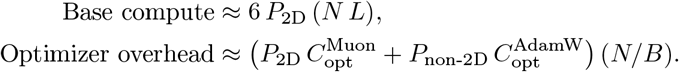

In our architecture *P*_non-2D_ ≪ *P*_2D_ (under 1% of total parameters), so we drop the non-2D overhead contribution and factor *P*_2D_ *N* out of the master formula. For brevity we write *P* for *P*_2D_ for the remainder of this appendix:

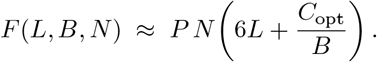

#### Per-optimizer step costs

Applying this to our four optimizers (with *C*_opt_ derived by counting scalar FLOPs per parameter for each kernel in the `step()` method of the PyTorch implementation, under our fused SwiGLU architecture):

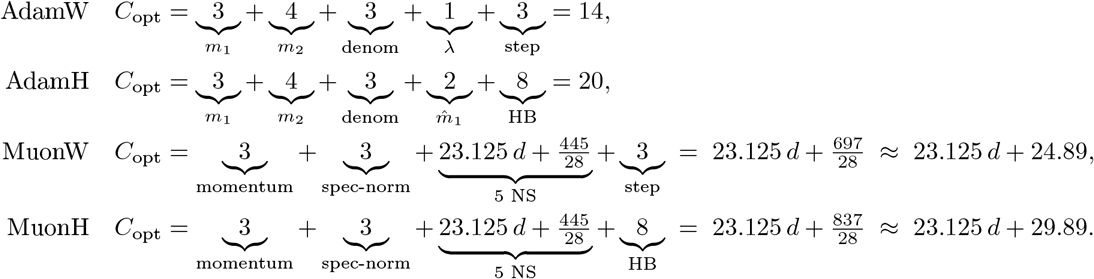

Each component counts the per-parameter FLOPs of one PyTorch kernel in the step (under the conventions *a*+=*αb* = 2 FLOPs/elt, `addcmul_ and addcdiv_` = 3 FLOPs/elt, and ∥*·* ∥_*F*_ = 2 FLOPs/elt). For Adam, *m*_1_ is the first-moment EMA (`mul_`+axpy), *m*_2_ is the second-moment EMA (`mul_ + addcmul_`), and “denom” is 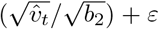 the AdamW tail is a decoupled weight-decay scale *λ* plus a fused *addcdiv_* “step” (1 + 3), whereas AdamH replaces it with a bias-correction divide 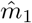 (`exp_avg`*/b*_1_, 2) and an 8-FLOP *Hyperball* (HB) projection {*u ← m/*denom, ∥*u*∥ _*F*_ , *u · R/* ∥*u*∥ _*F*_ , *p* += *ηu*, ∥*p*∥ _*F*_ , *p · R/*∥ *p*∥ _*F*_} . For Muon, the 3-FLOP “momentum” term is the in-place momentum-buffer update `buf.lerp_(grad, 1-m);` the 3-FLOP “spec-norm” is the Frobenius pre-divide Ũ *← U/* ∥*U*∥ _*F*_ that feeds Newton–Schulz; the MuonW/MuonH tails are identical to the AdamW/AdamH tails above (“step” = weight decay + scaled add, 1 + 2 = 3; “HB” = same 8-FLOP projection).

#### Deriving the Muon Newton–Schulz cost

The *d*-dependent 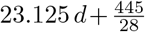 term sums the five Newton– Schulz iterations across all Muon-eligible weights in one transformer layer. Each iteration performs three matrix ops on a single weight of shape [*m, n*] with *m*≤ *n* (the code transposes if *m* > *n*): a Gram product *G* ← ŨŨ^⊤^ (2*m*^2^*n* matmul FLOPs), a cubic `addmm` *G* ← *bG* + *cG*^2^ (2*m*^3^ + 3*m*^2^), and a final `addmm` Ũ ← *a*Ũ + *G*Ũ (2*m*^2^*n* + 2*mn*). Per parameter, 5 iterations cost 20*m* + 10 *m*^2^*/n* matmul FLOPs and 10 + 15 *m/n* scaling FLOPs, so the cost depends on each weight’s post-transpose aspect ratio *r* = *m/n*. Summing over the six per-layer Muon-eligible weights of a grouped-query-attention (*g* ≡ num_kv_groups*/*num_heads = 2*/*8 = 1*/*4) + SwiGLU block with *d*_ff_ = 8*d/*3:

| Weight | $[m, n]$ | $r$ | Params | 5-step matmul | 5-step scaling |
| --- | --- | --- | --- | --- | --- |
| $W_q, \text{out\_proj}$ (each) | $[d, d]$ | 1 | $d^2$ | $30 d^3$ | $25 d^2$ |
| $W_k, W_v$ (each) | $[d/4, d]$ | $1/4$ | $d^2/4$ | $45 d^3/32$ | $55 d^2/16$ |
| `up_proj` | $[d, 16d/3]$ | $3/16$ | $16 d^2/3$ | $350 d^3/3$ | $205 d^2/3$ |
| `down_proj` | $[d, 8d/3]$ | $3/8$ | $8 d^2/3$ | $190 d^3/3$ | $125 d^2/3$ |
| **Per-layer total** | | | $10.5 d^2$ | $242.8125 d^3$ | $166.875 d^2$ |
| **Per parameter** | | | | $23.125 d$ | $\frac{445}{28} \approx 15.89$ |

These per-parameter averages are *exact* under the combination of grouped-query attention with *g* = 1*/*4 and *d*_ff_ = 8*d/*3 SwiGLU. For a uniformly-square set of weights the cost would be the cleaner 30*d* + 25; the [*d/*4, *d*] shape of *W*_*k*_, *W*_*v*_ and the 8*/*3 MLP expansion both pull the matmul coefficient down from 30*d* to 23.125 *d* and the constant from 25 to 445*/*28.

For our concrete architecture (*d* = 2048), MuonW’s and MuonH’s step costs evaluate to 47,384.89 and 47,389.89 FLOPs per parameter, respectively.

#### Checkpoint-to-FLOPs conversion

The FLOPs-axis plots map a checkpoint index *t* (a training step) to cumulative FLOPs via

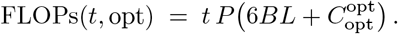

#### Overhead ratio

The fraction of compute consumed by the optimizer step relative to the base forward/backward pass is

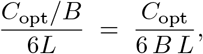

which is independent of the dataset size *N*. For our largest configuration (*L* = 352, *B* = 2400, *d* = 2048), MuonH’s *C*_opt_ ≈ 47,389.89 yields an overhead of roughly 0.94%, confirming that Muon’s Newton–Schulz orthogonalization step is a small fraction of total training FLOPs despite its *O*(*d*) scaling in *C*_opt_.

### G EMA Perplexity Values

The optimizer-level progression in Figure 5 selects one run per optimizer variant using the final exponential moving average (EMA) of checkpoint perplexities. The EMA decay is 0.9 and is applied over the sequence of checkpoint-level mean perplexities. This dampens batch-level and checkpoint-level noise: an isolated unusually easy or hard evaluation checkpoint changes the EMA by only 10% of its deviation from the previous smoothed value, while persistent improvements continue to accumulate across checkpoints. For Figure 5, the selected largest-scale runs are the minima in the hidden size (hs) 2048 columns below; these selections match the figure legend.

| AdamH |  |  |  |  |  |
| --- | --- | --- | --- | --- | --- |
| hs512 |  | hs1024 |  | hs2048 |  |
| LR | EMA PPL | LR | EMA PPL | LR | EMA PPL |
| 9.0e-05 | 2.6635 | 9.0e-05 | 2.6453 | 9.0e-05 | 2.6266 |
| 3.6e-04 | 2.6199 | 3.6e-04 | 2.6108 | 3.6e-04 | 2.5919 |
| 1.4e-03 | 2.5895 | 1.4e-03 | 2.5710 | 1.4e-03 | 2.5620 |
| 5.7e-03 | 2.5668 | 5.7e-03 | 2.5644 | 5.7e-03 | **2.5579** |
| 2.3e-02 | 2.5878 | 2.3e-02 | 2.5888 | 2.3e-02 | 2.5856 |

| AdamW |  |  |  |  |  |
| --- | --- | --- | --- | --- | --- |
| hs512 |  | hs1024 |  | hs2048 |  |
| LR | EMA PPL | LR | EMA PPL | LR | EMA PPL |
| 9.0e-05 | 2.6328 | 9.0e-05 | 2.6116 | 9.0e-05 | 2.5896 |
| 3.6e-04 | 2.5987 | 3.6e-04 | 2.5865 | 3.6e-04 | **2.5761** |
| 1.4e-03 | 2.5870 | 1.4e-03 | 2.5931 | 1.4e-03 | 2.5977 |
| 5.7e-03 | 2.5927 | 5.7e-03 | 2.5825 | 5.7e-03 | 2.5931 |
| 2.3e-02 | 2.5884 | 2.3e-02 | 2.5908 | 2.3e-02 | 2.5963 |

| MuonH |
| --- |

| hs512 |  | hs1024 |  | hs2048 |  |
| --- | --- | --- | --- | --- | --- |
| LR | EMA PPL | LR | EMA PPL | LR | EMA PPL |
| 9.0e-05 | 2.6496 | 9.0e-05 | 2.6306 | 9.0e-05 | 2.6156 |
| 3.6e-04 | 2.6096 | 3.6e-04 | 2.5993 | 3.6e-04 | 2.5841 |
| 1.4e-03 | 2.5837 | 1.4e-03 | 2.5675 | 1.4e-03 | 2.5538 |
| 5.7e-03 | 2.5654 | 5.7e-03 | 2.5476 | 5.7e-03 | **2.5474** |
| 2.3e-02 | 2.5957 | 2.3e-02 | 2.5924 | 2.3e-02 | 2.5952 |

MuonW
| hs512 |  | hs1024 |  | hs2048 |  |
| --- | --- | --- | --- | --- | --- |
| LR | EMA PPL | LR | EMA PPL | LR | EMA PPL |
| 9.0e-05 | 2.6902 | 9.0e-05 | 2.6818 | 9.0e-05 | 2.6721 |
| 3.6e-04 | 2.6335 | 3.6e-04 | 2.6262 | 3.6e-04 | 2.6161 |
| 1.4e-03 | 2.5987 | 1.4e-03 | 2.5848 | 1.4e-03 | 2.5724 |
| 5.7e-03 | 2.5708 | 5.7e-03 | 2.5492 | 5.7e-03 | 2.5359 |
| 2.3e-02 | 2.5550 | 2.3e-02 | 2.5345 | 2.3e-02 | **2.5306** |

SgdH
| hs512 |  | hs1024 |  | hs2048 |  |
| --- | --- | --- | --- | --- | --- |
| LR | EMA PPL | LR | EMA PPL | LR | EMA PPL |
| 9.0e-05 | 2.7320 | 9.0e-05 | 2.7202 | 9.0e-05 | 2.7151 |
| 3.6e-04 | 2.7088 | 3.6e-04 | 2.7011 | 3.6e-04 | 2.7035 |
| 1.4e-03 | 2.6903 | 1.4e-03 | 2.6881 | 1.4e-03 | 2.6724 |
| 5.7e-03 | **2.6402** | 5.7e-03 | 2.6467 | 5.7e-03 | 2.6907 |
| 2.3e-02 | 2.6894 | 2.3e-02 | 2.7326 | 2.3e-02 | 2.8241 |

SgdW
| hs512 |  | hs1024 |  | hs2048 |  |
| --- | --- | --- | --- | --- | --- |
| LR | EMA PPL | LR | EMA PPL | LR | EMA PPL |
| 9.0e-05 | 2.8186 | 9.0e-05 | 2.8071 | 9.0e-05 | 2.8024 |
| 3.6e-04 | 2.8022 | 3.6e-04 | 2.7965 | 3.6e-04 | 2.7914 |
| 1.4e-03 | 2.7894 | 1.4e-03 | 2.7809 | 1.4e-03 | 2.7839 |
| 5.7e-03 | 2.7788 | 5.7e-03 | 2.7733 | 5.7e-03 | 2.7743 |
| 2.3e-02 | **2.7694** | 2.3e-02 | 2.7921 | 2.3e-02 | 2.7877 |

### H Additional Figures

Table 1: Exact permutation tests (two-sided) for Pearson correlations between raw variables in *G. difficilis*, excluding Genovesa (considered an outlier). For each predictor–response pair, *r* is the Pearson correlation coefficient computed on untransformed data, and *p*_perm_ is the exact permutation *p*-value based on all 5! = 120 permutations under the null hypothesis of no association. Area and *N*_*e*_ are highly correlated. Data from (Lamichhaney et al., 2015).

## Footnotes

1 NerVE originally applies these metrics to track activation dynamics through FFN layers, but the same spectral measures are equally well defined for weight matrices.

